# T-lymphocyte Tyrosine Hydroxylase Regulates T_H_17 T-lymphocytes during Repeated Social Defeat Stress

**DOI:** 10.1101/2022.01.26.477910

**Authors:** Safwan K. Elkhatib, Cassandra M. Moshfegh, Gabrielle F. Watson, Adam J. Case

**Author notes:** **Corresponding author:** Adam J. Case, PhD, Associate Professor, Department of Psychiatry and Behavioral Sciences Department of Medical Physiology, College of Medicine, Texas A&M University, 3414 MREB2, 8447 Riverside Pkwy, Bryan, TX, 77807.

## Abstract

Posttraumatic stress disorder (PTSD) is a debilitating psychiatric disorder which results in deleterious changes to psychological and physical health. Patients with PTSD are especially susceptible to co-morbid inflammation-driven pathologies, such as autoimmunity, while also demonstrating increased T-helper 17 (T_H_17) lymphocyte-driven inflammation. While the exact mechanism of this increased inflammation is unknown, overactivity of the sympathetic nervous system is a hallmark of PTSD. Neurotransmitters of the sympathetic nervous system *(i.e.,* catecholamines) can alter T-lymphocyte function, which we have previously demonstrated to be partially mitochondrial redox-mediated. Furthermore, we have previously elucidated that T-lymphocytes generate their own catecholamines, and strong associations exist between tyrosine hydroxylase (TH; the rate-limiting enzyme in the synthesis of catecholamines) and pro-inflammatory interleukin 17A (IL-17A) expression within purified T-lymphocytes in a preclinical rodent model of PTSD. Therefore, we hypothesized that T-lymphocyte-generated catecholamines drive T_H_17 T-lymphocyte polarization through a mitochondrial superoxide-dependent mechanism during psychological trauma. To test this, T-lymphocyte-specific TH knockout mice (TH^T-KO^) were subjected to repeated social defeat stress (RSDS). RSDS characteristically increased tumor necrosis factor-α (TNFα), IL-6, IL-17A, and IL-22, however, IL-17A and IL-22 (T_H_17 produced cytokines) were selectively attenuated in circulation and in T-lymphocytes of TH^T-KO^ animals. When activated *ex vivo,* secretion of IL-17A and IL-22 by TH^T-KO^ T-lymphocytes was also found to be reduced, but could be partially rescued with supplementation of norepinephrine specifically. Interestingly, TH^T-KO^ T-lymphocytes were still able to polarize to T_H_17 under exogenous polarizing conditions. Last, contrary to our hypothesis, we found RSDS-exposed TH^T-KO^ T-lymphocytes still displayed elevated mitochondrial superoxide, suggesting increased mitochondrial superoxide is upstream of T-lymphocyte TH induction, activity, and T_H_17 regulation. Overall, these data demonstrate TH in T-lymphocytes plays a critical role in RSDS-induced T_H_17 T-lymphocytes and offer a previously undescribed regulator of inflammation in RSDS.

## Introduction

Posttraumatic stress disorder (PTSD) is a debilitating, stress-related disorder which has been described by various names since antiquity. PTSD is characterized by exposure to a traumatic event in conjunction with the persistence of a constellation of symptom clusters meeting specific diagnostic and statistical manual of mental disorders (DSM) criteria, such as affective changes, avoidance behavior, dissociative symptoms, and hyperarousal.^1^ PTSD as a mental health disorder presents a tremendous cost to the individual and society, with 40% of patients suffering from PTSD even 10 years after their initial diagnosis.^2, 3^

Importantly, there is a breadth of research demonstrating a connection between PTSD and deleterious changes to physical health.^4^ Patients with PTSD are at an increased risk for a variety of inflammation-driven diseases, ranging from rheumatoid arthritis to cardiovascular disease,^5–8^ which ultimately contribute to the decreased quality of life and lifespan of these patients. Intimately related to this risk for co-morbid diseases, patients with PTSD display distinctive physiological changes, specifically within the nervous and immune systems. Patients with PTSD demonstrate increased sympathetic tone by measures such as urinary norepinephrine (NE) and baroreflex sensitivity,^9–11^ in addition to heightened T-lymphocyte-driven inflammatory markers.^12–14^ There is a breadth of work demonstrating the effects of NE on T-lymphocyte functional status both *in vitro* and *in vivo*.^15^ For example, our *in vitro* work demonstrated NE supplementation to activated T-lymphocytes resulted in increased pro-inflammatory cytokine profiles, such as IL-6 and IL-17A, partially mediated by a mitochondrial redox mechanism.^16^ Thus, the link between PTSD and inflammatory diseases may lie in connection of catecholamines and T-lymphocytes.

Critically, it has been shown in numerous investigations that T-lymphocytes themselves generate, release, and respond catecholamines (dopamine, norepinephrine, and epinephrine). ^18^ This has been demonstrated at the single cell level in both isolated and immortalized lymphocytes, while *in vitro* studies utilizing pharmacological inhibition of catecholamine synthesis demonstrated these catecholamines influence the functionality and activation state of T-lymphocytes.^18^ In our own work, we investigated how increased sympathoexcitation in the murine model of PTSD known as repeated social defeat stress (RSDS) could affect T-lymphocyte inflammatory signatures.^19^ We discovered that T-lymphocytes from RSDS animals had increased in the gene expression of tyrosine hydroxylase (TH; the rate-limiting step in catecholamine synthesis), which strongly correlated with expression of pro-inflammatory IL-17A in purified T-lymphocytes. Importantly, IL-17 and T_H_17 cells have been heavily implicated in several inflammatory and autoimmune diseases.^20, 21^

To this end, we sought to examine the role of T-lymphocyte-generated catecholamines in T-lymphocyte-driven inflammation seen during RSDS. By generating a conditional T-lymphocyte TH knockout mouse (TH^T-KO^), we were able to selectively investigate this mechanism and further define the role for catecholamines in T_H_17-mediated inflammation.

## Materials and Methods

### Mice

All experimental mice were bred inhouse in a room separate from stress induction and stress-exposed mice to reduce physical, psychological, and social stressors. Littermates were group housed (≤5 mice per cage) prior to RSDS or control protocol initiation at 8-12 weeks of age. All mice were housed with standard corncob bedding, paper nesting material, and given access to standard chow (Teklad Laboratory Diet #7012, Harlan Laboratories, Madison, WI) and water ad libitum. At the completion of experiments, experimental mice were euthanized by pentobarbital overdose (150 mg/kg, Fatal Plus, Vortech Pharmaceuticals, Dearborn, MI) administered by intraperitoneal injection. Daily RSDS and euthanasia occurred between 0700 and 1000 Central Standard Time to attenuate circadian rhythm effects on the neuroendocrine and immune systems.

In order to specifically investigate the role of TH in T-lymphocytes, a cell-type specific (conditional) TH knockout mouse was created and utilized herein. First, mice possessing loxP elements flanking exon 1 of the TH gene locus *(i.e.,* B6.Cg-Th^tm4.1-Rpa^; TH^lox/lox^) were graciously obtained from Michael Iuvone at Emory University.^22^ TH^lox/lox^ mice were then crossed to mice expressing codon-improved cre recombinase under the control of the distal promoter of T-lymphocyte-specific tyrosine kinase (Lck) *(i.e.,* B6.Cg-Tg(Lck-icre)^3779Nik/J^), originally generated by Nigel Killeen.^23^ This distal Lck promoter is active at or after T-cell receptor (TCR) upregulation during positive selection of thymocytes and has shown activity in both αδ and γδ T-lymphocyte subsets.^23^ Mice were crossed to the F3 generation, allowing for 100% progeny possessing homozygous TH^lox/lox^ alleles, 50% bearing Lck cre recombinase (**Figure 1A**; T-lymphocyte specific TH knockouts; referred to as TH^T-KO^), and 50% cre negative (**Figure 1A**; TH^lox/lox^ cre recombinase negative controls; referred to as TH^Con^).

**Figure 1.**
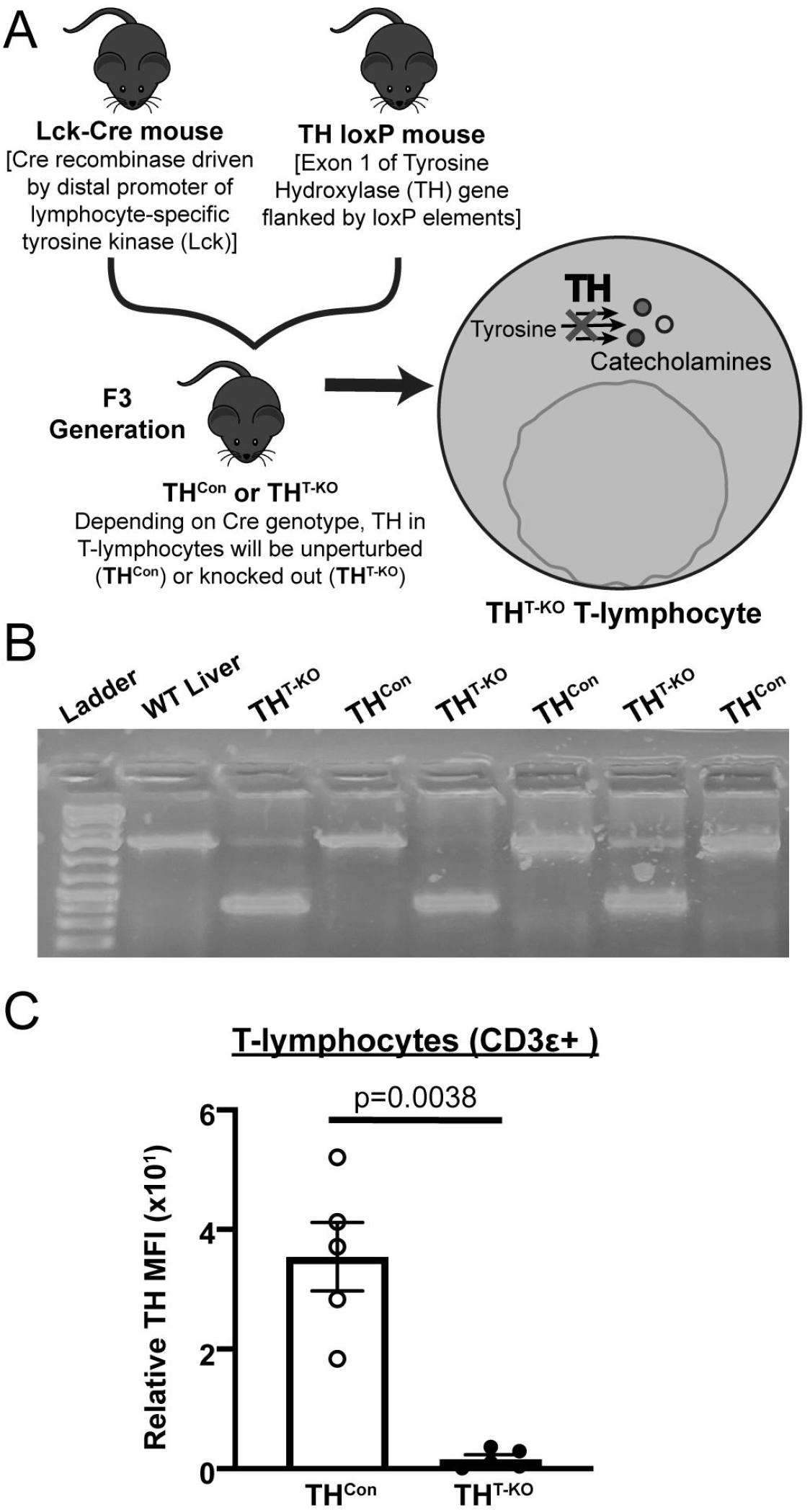
Generation and validation of a conditional T-lymphocyte TH knockout model. A) Through crossing of TH loxP and Lck-cre mice to the F3 generation, 50% conditional genetic knockout (TH^T-KO^) and 50% cre-negative (TH^Con^) mice were generated, allowing for *in vivo* investigation of the role of TH within T-lymphocytes was generated. B) Representative DNA gel of respective samples following PCR amplification with oligonucleotide primers which produced differentially sized PCR products. C) Quantification of flow cytometry of TH mean fluorescence intensity (MFI) in splenic T-lymphocytes (CD3ε+) in TH^T-KO^ and TH^Con^; significance by Mann-Whitney U test, all values normalized to TH FMO tube.

When possible, experimenters were blinded to the genotype and stress-exposure of mice until the completion of the study. Care was taken to blind experimenters to conditions during biological assay and data acquisition. All procedures were reviewed and approved by the University of Nebraska Medical Center and Texas A&M University Institutional Animal Care and Use Committees.

### Genotyping TH KO

Rationally designed primers and polymerase chain reaction (PCR) were utilized to assess recombination of the TH gene. DNA was first isolated through use of GeneJET Genomic DNA Purification Kit (#K0722, Thermo scientific, Waltham, MA). Oligonucleotide primers (Forward: 5’-GAAGACCCTAGGGAGATGCCAAA-3’; Reverse 5’ TTTCCCTTACTTCA-CAAAATAGGACCCACAGAA) were specifically designed which could distinguish the recombined KO TH allele from the unperturbed TH^lox/lox^ genotype by size. PCR products were loaded onto a 1.5% agarose gel with 1 mg/mL of ethidium bromide for intercalation, separated by electrophoresis at 100 V, and visualized under ultraviolet light alongside a DNA ladder (GeneRuler 1kb Plus, #SM1331, Thermo Scientific).

### Repeated social defeat stress paradigm

An adapted version of RSDS was performed as we have previous described.^19, 24, 25^ Briefly, RSDS exposes experimental mice to psychosocial trauma through repeated daily interaction and subsequent shared housing with 4–8-month-old aggressive, retired breeder mice of a CD-1 background (Charles River #022, Wilmington, MA). Due to usage of retired male breeders for stress induction, female mice and sex differences are precluded from study by conventional RSDS. During the 10-day RSDS paradigm, each experimental mouse was placed into the home cage of a CD-1 mouse for five minutes daily to allow for physical confrontation, while control mice remained in home cages. For the remaining 24 hours, experimental and CD-1 mice were co-housed within the same cage but separated by a transparent, perforated barrier. This paradigm was repeated daily by rotating the experimental mouse to a novel CD-1 mouse and its respective cage, while control mice remained pair housed with other control mice throughout the duration of the RSDS paradigm. Mice were promptly excluded from further study if they demonstrated evident signs of wounding (>1 cm) or lameness during or after RSDS. At the end of the 10-day period (day 11), all mice underwent behavioral testing followed by euthanasia and tissue harvest the following day (day 12).

### Elevated Zero Maze

In order to assess anxiety-like behavior, the elevated zero maze test was utilized as has been previously described^19^. Briefly, the maze is comprised of a raised circular platform maze divided into quadrants, with two open portions and two closed arms. Following RSDS or control exposure, mice explore the novel environment for five minutes, with each session recorded and subsequently analyzed using Noldus Ethovision XT13 software (Noldus Information Technology).

### Social Interaction Test

Following RSDS or control housing, pro-social and depressive-like behavior was assessed by the social interaction test as has been previously described.^19^ Briefly, each experimental mouse was placed within an open field chamber containing a small mesh enclosure in two separate 2.5-minute sessions. In the first session, all mice were tested with an empty enclosure. Next, a novel CD-1 mouse was placed within the enclosure and experimental mice were again recorded. All sessions were recorded, tracked, and analyzed with Noldus Ethovision XT13 software.

### Splenic T-lymphocyte isolation

Pan, CD4+, and CD8+ murine T-lymphocytes were magnetically negatively selected from whole spleens as has been previously described.^16^ Briefly, murine spleens were disassociated by ground glass slides, resuspended in supplemented RPMI media, then passed through a 70 μM filter. Splenic T-lymphocytes were isolated using EasySep Mouse T-cell Isolation Kit (#19851, STEMCELL™) per manufacturer’s instructions.

### T-lymphocyte activation, culture, and catecholamine supplementation

T-lymphocytes were cultured as has been previously described.^26^ Briefly, after T-lymphocyte isolation, 8.0 x 10^5^ cells/mL were plated with anti-CD3/anti-CD28 magnetic activation beads in a 1:1 ratio with cells (Dynabeads™, #11456D, Thermo Fisher Scientific), and incubated for 72 hours at 37°C prior to harvest and analysis.

For catecholamine treatment conditions, freshly harvested splenic CD4+ T-lymphocytes were isolated and plated in 24 well plates. Wells were supplemented daily with 10 μM of dopamine hydrochloride #AAA1113606, Thermo Scientific Chemicals), L-norepinephrine (#AAL0808703, Thermo Scientific Chemicals), or L-epinephrine (#AAL0491106, Thermo Scientific Chemicals) reconstituted in 1x sterile PBS. Catecholamine concentrations were based on our previous published work with *ex vivo* catecholamine dosage curves of T-lymphocytes.^16,27^ T-lymphocytes from a single animal received each treatment condition allowing for paired analyses, with 3 technical replicates for each sample and treatment combination.

T-lymphocyte growth curves were obtained by live cell analyses. CD4+ or CD8+ T-lymphocytes were isolated and plated as stated above in 96-well flat bottom plates and then placed into an Incucyte S3 Live-cell analysis system (Essen Bioscience) housed within an environmental chamber (37°C, 5% CO_2_) for 72 hours. Live cell images for confluence assessments were taken every 5 minutes for the 72-hour period. Confluence was normalized to starting densities after optical focusing, with >5 technical replicate wells used per sample. Data were analyzed using Incucyte S3 analysis software.

### T-lymphocyte T_H_17 Polarization

CD4+ splenic T-lymphocytes were polarized to T_H_17 *ex vivo* by use of CellXVivo Mouse Th17 Cell Differentiation Kit (#CDK017, R&D Systems). Briefly, CD4+ T-lymphocytes were isolated and cultured as described above with RPMI media additionally supplemented with a cocktail of proprietary polarizing reagent antibodies which promote T_H_17 polarization and prevent T_H_1 and T_H_2 differentiation. Following 5 days of culture and activation, T-lymphocytes were harvested for flow cytometric staining and analysis.

### Flow cytometric cellular staining

Extracellular and intracellular flow cytometric staining for T-lymphocyte populations was performed as has been previously described.^25^ Briefly, cells (were incubated at 37° C for 4 hours in RPMI media supplemented with phorbol 12-myristate-13-acetate (PMA; 10 ng/mL), ionomycin (0.5 mg/mL), and BD GolgiPlug Protein Transport Inhibitor (containing brefeldin A; 1 mg/mL; BD Biosciences). Cells were washed, resuspended in PBS, and amine-reactive viability stained for 30 minutes at 4° C with Live/Dead Fixable Cell Stain Kit (#L34960, Thermo Fisher Scientific). Next, cells were washed and resuspended in RPMI media supplemented with antibodies targeting the following extracellular markers: CD3ε PE-Cy7 (clone 145-2C11, eBioscience), CD4 Alexa Fluor 488 (clone GK1.5, eBioscience), and CD8a APC (clone 53-6.7, BD Biosciences). In order to stain for intracellular proteins, cells were then fixed and permeabilized utilizing the FOXP3 Fixation and Permeabilization kit (#00-5523-00, eBioscience) per the manufacturer’s instructions. Cells were washed again and resuspended for 30 minutes in permeabilization buffer containing IL-17A PE (clone TC11-18H10, BD Biosciences) and primary rabbit anti-tyrosine hydroxylase antibody (clone EP1533Y, Abcam). Next, cells were washed and resuspended in RPMI media supplemented with secondary goat anti-rabbit QDOT 605 (Q11402MP, Invitrogen). Cells were then washed and resuspended in cold PBS for immediate analysis. Data were acquired using a customized BD LSRII flow cytometer (UNMC, BD Biosciences) or a 4-laser Attune NxT flow cytometer (Thermo Fisher Scientific). All flow cytometry experiments were completed with single-color and fluorescence minus one (FMO) control samples, with indirect intracellular staining also employing a primary antibody only control sample. All analyses were conducted on FlowJo 10 software (BD Bioscience).

### Flow cytometric redox assessment

Mitochondrial-specific assessment of superoxide was performed as has been previously described.^16^ Briefly, cells were stained with cell-type specific fluorescent antibodies [anti-CD3ε PE-Cy7 (clone 145-2C11, eBioscience), CD19 APC-Cy7 (clone 6D5, BioLegend), CD11b SB-436 (clone M1/70, eBioscience), CD11c APC (clone N418, eBioscience), and NK1.1 SB-600 (clone PK136, eBioscience)] in addition to 1 μM of O_2_^•-^-sensitive mitochondrial-localized probe, MitoSOX Red (#M36008, Thermo Fisher Scientific) for 30 minutes at 37°C. Cells were analyzed on an LSRII flow cytometer at 488/610 nm excitation/emission, respectively, and data analyzed using FlowJo software.

### RNA extraction, cDNA production, and real-time RT-qPCR

Gene expression assessment of purified T-lymphocytes was performed as previously described.^19^ Briefly, mRNA was extracted from purified T-lymphocytes by silica-membrane spin columns through use of RNAeasy mini kit (#74104, Qiagen), then immediately converted to cDNA by High-Capacity cDNA Reverse Transcription Kit (#4374966, Applied Biosystems). Generated cDNA was used for real-time qPCR using respective gene targeted intron-spanning gene-specific unlabeled primers and dual-labeled fluorescent FAM probes with a proprietary quencher molecule (PrimePCR™, Bio-Rad, Hercules, CA). Thresholds were set objectively to determine cycle thresholds (CT), with 40S ribosomal protein S18 (RPS18) utilized as a loading control to determine ΔCT. All values were normalized to TH^Con^ control samples to determine Δ△CT values, which were then transformed to generate fold changes by the 2^-ΔΔCT^ method.

### Cytokine analyses

Plasma and spent culture media were harvested and analyzed for cytokine content by electrochemiluminescence as has been previously described.^24^ Briefly, blood was harvested by cardiac puncture immediately following sacrifice and anticoagulated with EDTA. Plasma was separated by centrifugation and stored at −80°C until assay. Spent media was harvested after 72 hours of T-lymphocyte activation. First, antigenic activation beads were magnetically removed, followed by centrifugation at 500xg for 2 minutes to pellet T-lymphocytes for downstream applications. Supernatant was removed and stored at −80°C until assay. Cytokine analyses were conducted through use of U-PLEX T_H_17 Meso Scale Discovery Kit (#K15078K, Meso Scale Discovery). All cytokine analyses were conducted per manufacturer’s instructions and quantified on a Meso Scale Discovery Quickplex SQ 120, with analyses conducted using Mesoscale Discovery Workbench software.

### Statistics

*A* total of 107 animals (53 TH^Con^, 54 TH^T-KO^) were utilized for the studies described herein. All data are presented as mean ± standard error of the mean (SEM) with sample numbers displayed as individual markers, same for repeated measures experiments, where n values are included within figure legend. Due to the nature of the experiments herein, not every biological assay was completed on each mouse within the RSDS or control paradigms. At least 3 independent experimental repeats were conducted for each experimental design. For comparisons with two independent groups, Shapiro-Wilk normality was performed followed by statistical testing by Mann-Whitney U or Student t-test as appropriate. In experiments with two levels of categorical variables (such as RSDS/Control, and TH^Con^/TH^T-KO^), full model two-way ANOVA was utilized. Two-way ANOVA Šidák multiple comparison tests of interest are listed in figures if respective group or interaction effects were found to be significant, with ANOVA p values listed within figure legend. All statistics were completed in GraphPad Prism (V9, GraphPad).

## Results

### TH^T-KO^ mice are viable model to assess TH within T-lymphocytes

In order to investigate the role of TH in T-lymphocytes exclusively, a conditional knockout mouse was generated and validated (**Figure 1A**). In Lck-driven cre-expressing cells *(i.e.,* T-lymphocytes), exon 1 of the TH gene was successfully removed, resulting in an ~800 bp shifted product by PCR (**Figure 1B**). Due to the likely imperfect T-lymphocyte purification, faint non-recombined bands can be seen within TH^T-KO^ T-lymphocyte PCR products (**Figure 1B**). Furthermore, TH^Con^ and TH^T-KO^ flow cytometric assessment demonstrated nearly absent TH within T-lymphocytes [CD3ε+; p=0.0038, **Figure 1C**). Overall, these data demonstrate effective knockout of TH within TH^T-KO^ Pan T-lymphocytes.

### Behavioral responses to RSDS are not altered in TH^T-KO^ animals

Following exposure to RSDS or control paradigms, TH^Con^ and TH^T-KO^ were assessed for differences in sociability and anxiety-like behavior by social interaction test and elevated zero maze, respectively. By elevated zero maze, control TH^T-KO^ demonstrated less distance moved after RSDS-exposure, independent of T-lymphocyte TH (p=0.0059, TH^Con^ Con vs. RSDS, and p=0.0067, TH^T-KO^ Con vs. RSDS; **Figure 2A**). Time within the open arm was significantly different by RSDS only (p=0.0040, Two-way ANOVA), with multiple comparison tests nearing significance (p=0.1114, TH^Con^ Con vs. RSDS, and p=0.1114, TH^T-KO^ Con vs. RSDS; **Figure 2A**). From the social interaction ratio, RSDS-exposed TH^Con^ and TH^T-KO^ mice demonstrated significant reductions in sociability (p=0.0063, TH^Con^ Con vs. RSDS, and p=0.0159, TH^T-KO^ Con vs. RSDS; **Figure 2B**), with no effect of genotype (p=0.9233, Two-way ANOVA). These assessments demonstrate established behavioral changes following RSDS exposure, with TH T-lymphocyte having no influence on these RSDS-induced behaviors.

**Figure 2.**
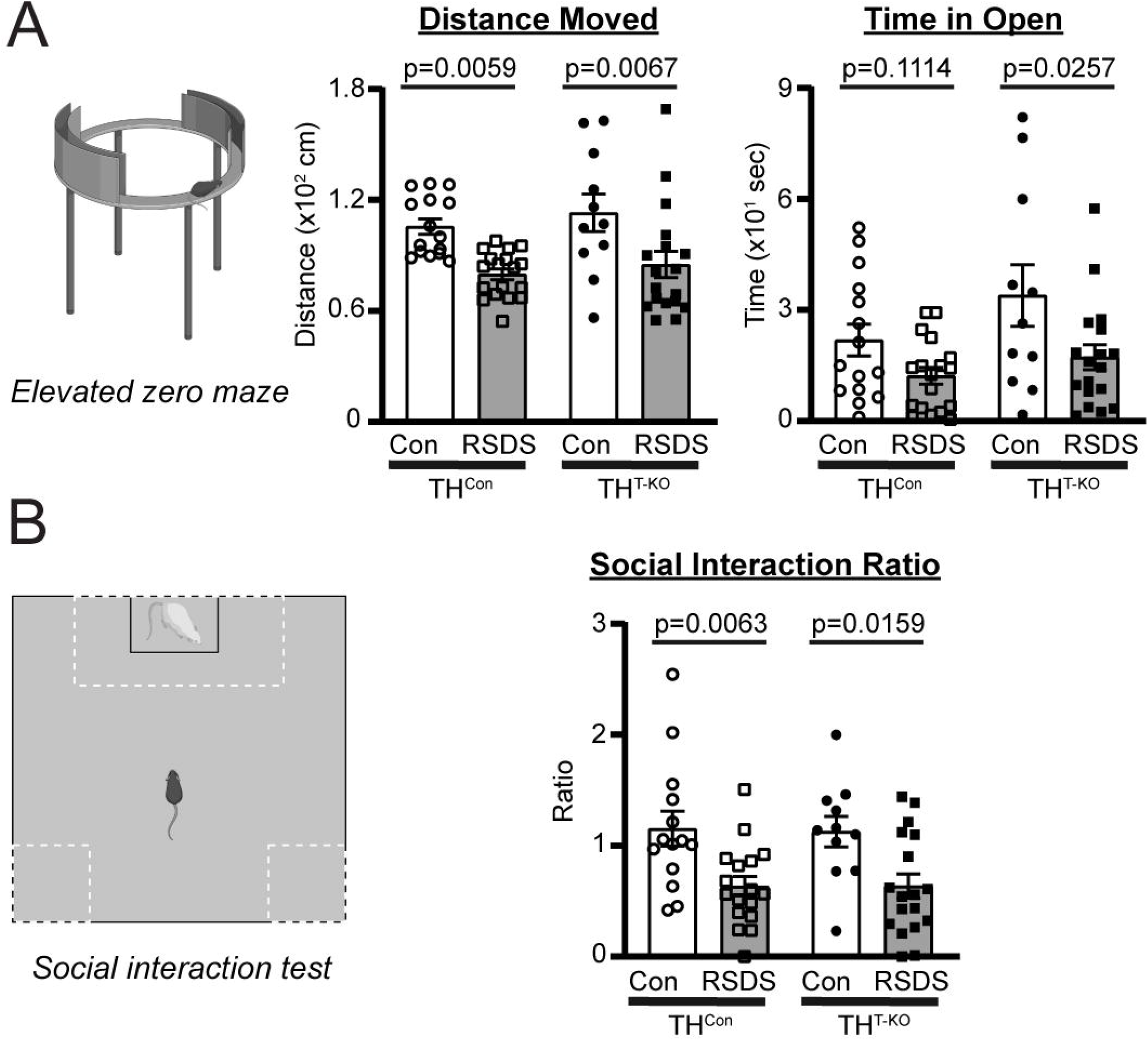
T-lymphocyte TH knockout does not affect RSDS-induced anxiety-like or social behavior. Mice were tested for anxiety-like and pro-social behavior following control or RSDS paradigms. A) *Left,* Representation of elevated zero maze for anxiety-like behavior. *Middle,* Total distance moved on elevated zero maze, Two-way ANOVA results; stress p<0.0001, genotype p=0.3042, interaction p=0.8672. *Right,* Time in open arm of elevated zero maze, Two-way ANOVA results; stress p=0.0040, genotype p=0.0575, interaction p=0.4305. B) *Left,* Representation of social interaction test for social behavior. *Right,* Social interaction ratio (defined as time spent with CD-1 present/time spent with empty enclosure), Two-way ANOVA results; stress p=0.0001, genotype p=0.9233, interaction p=0.9101. Two-way ANOVA Šidák multiple comparison tests of interest are listed if respective group or interaction effects were found to be significant.

### Circulating T_H_17 cytokines differ in TH^T-KO^ mice after RSDS, reflecting changes in T_H_17 populations and cytokine expression

We have previously demonstrated RSDS significantly increases circulating pro-inflammatory cytokines in WT mice, such as IL-2, IL-6, IL-10, IL-17A, IL-22, TNFα, CCL2, and CXCL2^24^. Importantly, many of these cytokines are related to adaptive immune function and can be generated by T-lymphocytes. In particular, IL-17A and IL-22 are cytokines released in large quantities by T_H_17 T-lymphocytes, a subtype of CD4+ T-lymphocytes which have been implicated in autoimmunity.^20^ In TH^Con^ animals, RSDS induced a significant increase in many of these cytokines paralleling earlier studies, such as IL-6, TNFα, IL-17A, and IL-22, despite smaller sample sizes across the four groups (p=0.0429, 0.0004, 0.0003, and 0.0054, respectively, TH^Con^ Con vs. RSDS; **Figure 3A**). In TH^T-KO^ mice, IL-6 and TNFα in circulation were increased after RSDS (p=0.0571 and <0.0001, respectively, TH^T-KO^ Con vs. RSDS; **Figure 3A**). However, this increase in RSDS-exposed TH^T-KO^ animals was significantly attenuated in IL-17A and IL-22, specifically (p=0.8930 and 0.9994, respectively, TH^T-KO^ Con vs. RSDS; **Figure 3A**), with differences in IL-17A and IL-22 by genotype in RSDS-exposed animals (p=0.0058 and 0.0061, respectively, TH^Con^ RSDS vs. TH^T-KO^ RSDS; **Figure 3A**).

**Figure 3.**
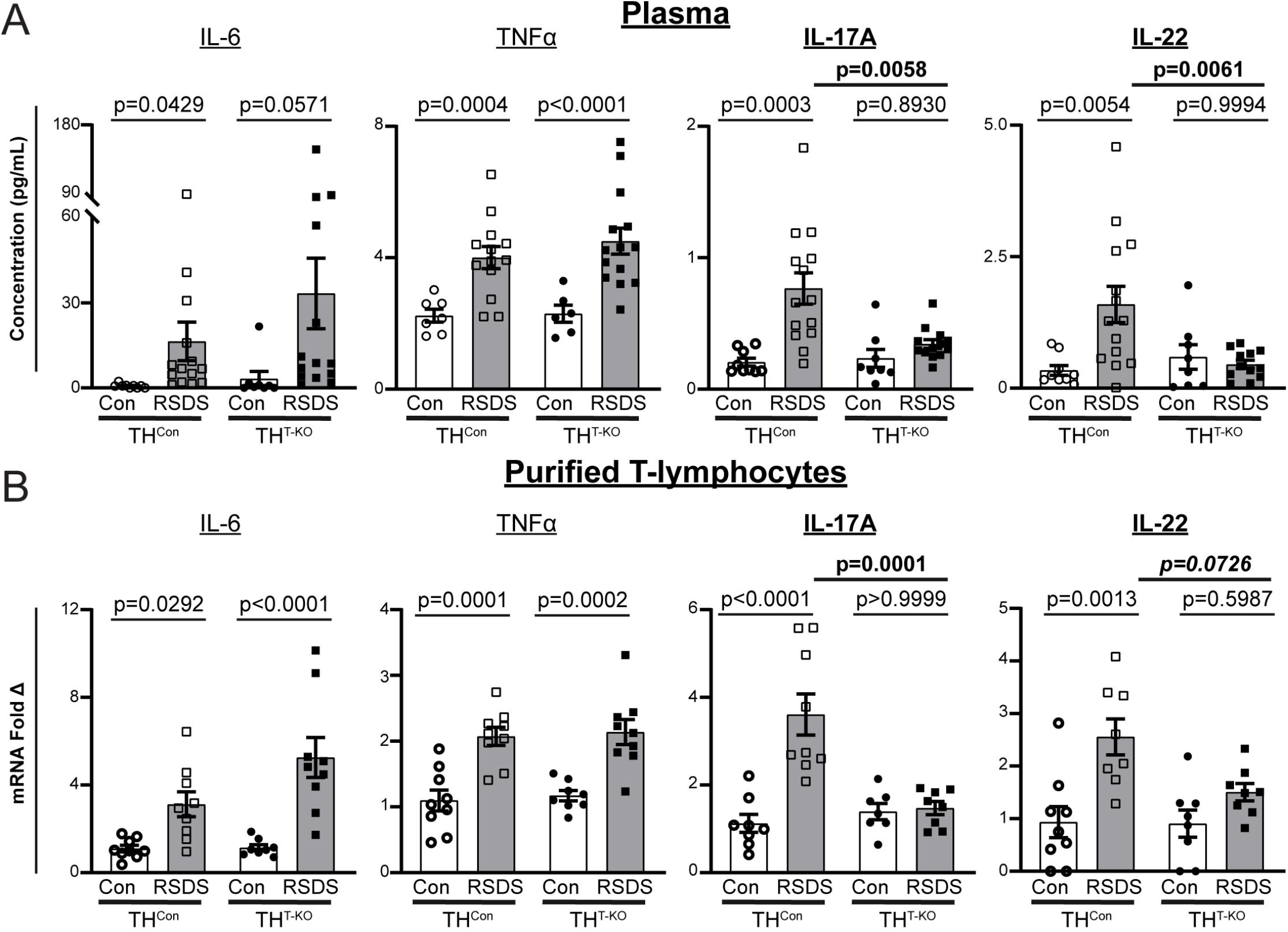
T-lymphocyte TH knockout selectively attenuates IL-17A and IL-22 cytokine levels. A) Circulating cytokines were assessed following control or RSDS paradigms with Meso Scale Discovery T_H_17 (Combo 2) assay. IL-6 Two-way ANOVA results, Stress p=0.0173, genotype p=0.2997, interaction 0.4460; TNFα Two-way ANOVA results, Stress p<0.0001, genotype p=0.5505, interaction 0.513; IL-17A Twoway ANOVA results, Stress p=0.0004, genotype p=0.0473, interaction p=0.0222; IL-22 Two-way ANOVA results, Stress p=0.0352, genotype p=0.0908, interaction 0.0094. B) Splenic T lymphocytes were isolated by magnetic negative selection, followed by RNA extraction, cDNA conversion, and real-time qPCR. All ΔCT values normalized to 40S ribosomal protein S18 (RPS18) control, then fold change normalized to TH^Con^ Control animals. IL-6 Two-way ANOVA results, Stress p<0.0001, genotype p=0.0619, interaction 0.0733; TNFα Two-way ANOVA results, Stress p<0.0001, genotype p=0.6486, interaction 0.9768; IL-17A Two-way ANOVA results, Stress p=0.0003, genotype p=0.0051, interaction p=0.0005; IL-22 Two-way ANOVA results, Stress p=0.0004, genotype p=0.0735, interaction 0.0593. Two-way ANOVA Šidák multiple comparison tests of interest are listed if respective group or interaction effects were found to be significant.

To further investigate these cytokine changes, splenic T-lymphocytes were isolated for gene expression analyses by real-time qPCR. We observed statistically significant increases in IL-6 and TNFα gene expression independent of genotype (p=0.0292 and 0.0001, respectively, TH^Con^ Con vs. RSDS; p<0.0001 and 0.0002, respectively, TH^T-KO^ Con vs. RSDS; **Figure 3B**). However, we demonstrated a similar attenuation of RSDS-induced increases in expression of IL-17A, with a trend toward significance in IL-22, in TH^T-KO^ T-lymphocytes as compared to TH^Con^ (p=0.0001 and 0.0726, respectively, TH^Con^ RSDS vs. TH^T-KO^ RSDS; **Figure 3B**).

We next investigated T_H_17 populations in our TH^T-KO^ and RSDS paradigms by a multi-parametric flow cytometric panel to identify T_H_17 T-lymphocytes (CD3ε+CD4+CD8-IL-17A+ splenocytes; **Figure 4**). RSDS induced an increase in T_H_17 T-lymphocytes in TH^Con^ mice but had had no effect T_H_17 populations of TH^T-KO^ mice (p=0.0326, TH^Con^ Con vs. RSDS; p>0.9999; TH^T-KO^ Con vs. RSDS; **Figure 4**), with significant differences in RSDS-exposed animals by genotype (p=0.0032, TH^Con^ RSDS vs. TH^T-KO^ RSDS; **Figure 4**). Overall, these data demonstrate RSDS induces T_H_17 polarization and cytokine production in a TH-dependent fashion within T-lymphocytes.

**Figure 4.**
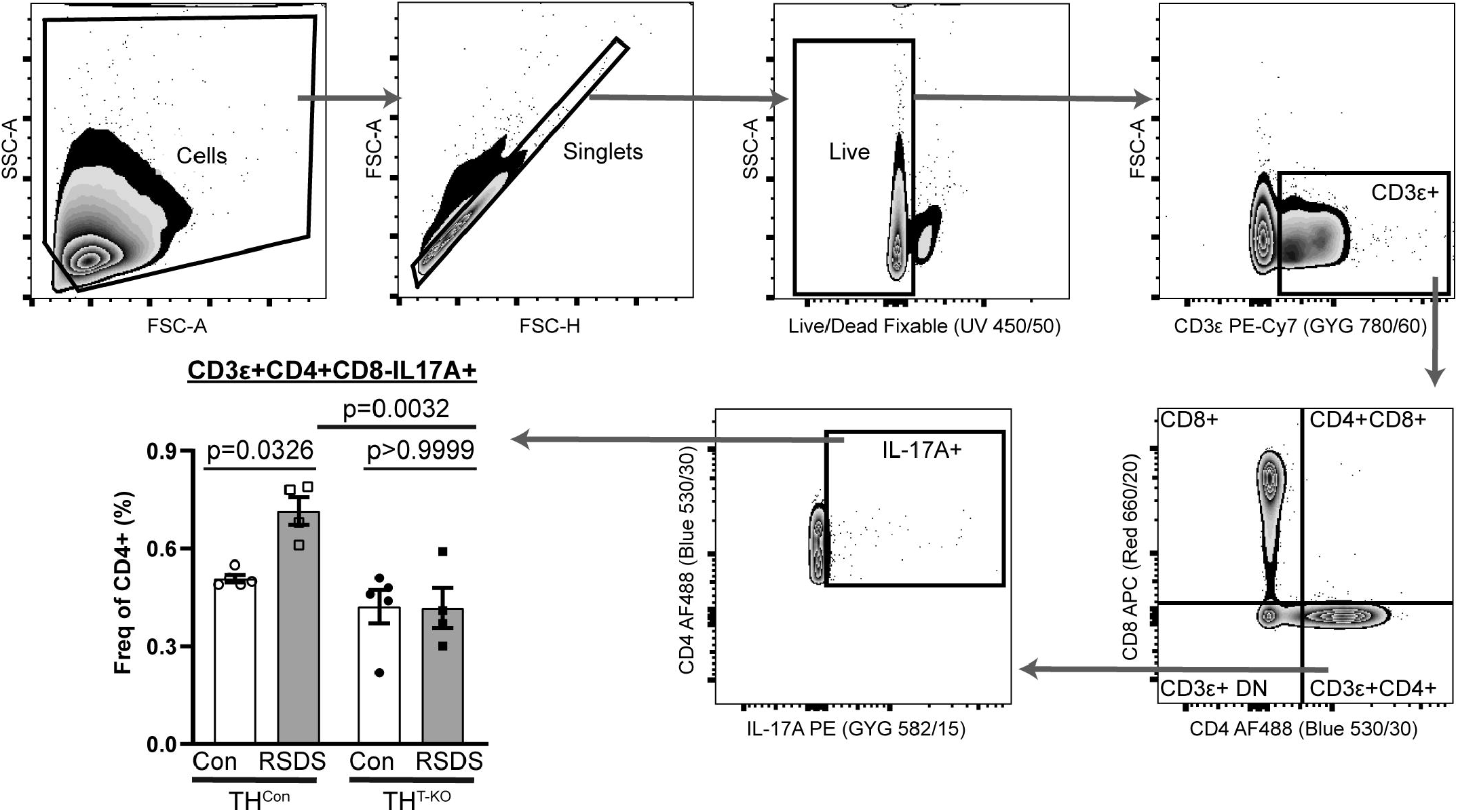
TH animals have attenuated T_H_17 polarization following RSDS. Following RSDS or control paradigms, splenocytes were harvested for multiparametric flow cytometry for T_H_17 T-lymphocytes (CD3ε+CD4+CD8-IL-17A+). Hierarchical gating strategy was employed to assess viability and percentage T_H_17 positivity. Two-way ANOVA results, Stress p=0.0398, genotype p=0.0007, interaction 0.0328. Two-way ANOVA Šidák multiple comparison tests of interest are listed if respective group or interaction effects were found to be significant.

### TH^T-KO^ T-lymphocytes demonstrate altered secretion of IL-17A and IL-22 despite similar growth

To focus our investigation on the role of TH in T-lymphocytes, TH^T-KO^ CD4+ and CD8+ T-lymphocytes from control (unstressed) animals were isolated for culture and activated by antigen-independent anti-CD3/CD28 stimulatory beads. Utilizing live-cell imaging, we detected no differences in growth in TH^T-KO^ CD4+ or CD8+ T-lymphocytes (p=0.6912 and 0.3516, respectively; **Figure 5A**). After 72 hours of activation, cytokine analyses of spent media of TH^T-KO^ CD4+ T-lymphocytes revealed altered content of critical pro-inflammatory cytokines; IL-6 and TNFα were unchanged or trended towards statistical significance (p=0.8353 and 0.0606, respectively; **Figure 5B**) while IL-17A and IL-22 were significantly decreased (p=0.0040 and 0.008, respectively; **Figure 5B**).

**Figure 5.**
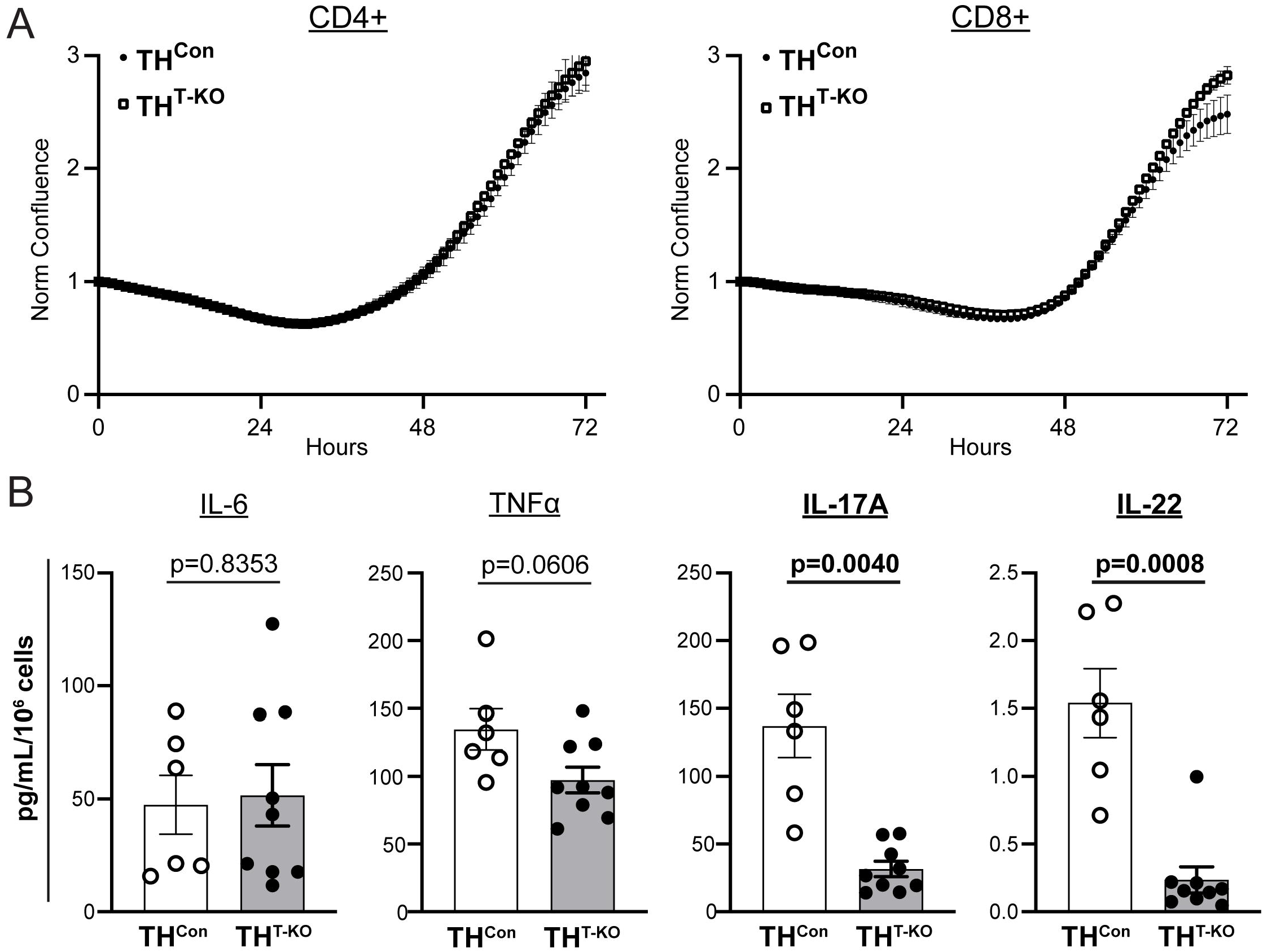
TH^T-KO^ CD4+ T-lymphocytes demonstrate altered T_H_17 cytokine secretion, but similar growth. A) CD4+ or CD8+ T-lymphocytes were isolated by negative selection, plated with anti-CD3/CD28 beads, and imaged for confluence 72 hours. N=4 samples, with >5 technical replicates per well. CD4+ Repeated measures Two-way ANOVA results, Genotype p=0.6912, Time p<0.0001, Interaction p>0.9999; CD8+ Repeated measures Two-way ANOVA results, Genotype p=0.3516, Time p=0.0003, Interaction p=0.0002. B) CD4+ T-lymphocytes were isolated and activated as above, with spent culture media collected and assessed for cytokines by Meso Scale Discovery T_H_17 (Combo 2) assay, with concentrations normalized to final cell counts. Listed p values were calculated by unpaired t-test or Mann-Whitney U test where appropriate from Shapiro-Wilk normality testing.

These cytokines were found unchanged or undetectable in CD8+ T-lymphocyte culture media (data not shown). Overall, this investigation provides further evidence of differences in TH^T-KO^ T-lymphocytes ability to produce T_H_17 cytokines.

### TH^T-KO^ T-lymphocyte IL-17A and IL-22 cytokine production can be restored with catecholamine supplementation

To explore the role of specific catecholamines in TH^Con^ and TH^T-KO^ CD4+ T-lymphocyte cytokine production, CD4+ T-lymphocytes were isolated, activated, and cultured *ex vivo* with 10 μM of each dopamine, norepinephrine, or epinephrine for paired analyses after 72 hours. IL-17A and IL-22 production was rescued primarily by NE supplementation (p=0.7989, TH^Con^ Vehicle vs. TH^T-KO^ NE; **Figure 6A**), with dopamine and epinephrine supplementation demonstrating modest effects. TNFα demonstrated no significant changes with catecholamine supplementation or by genotype (**Figure 6A**). Overall, this further validates a role for T-lymphocyte TH and subsequent catecholamines in T_H_17 salient cytokines.

**Figure 6.**
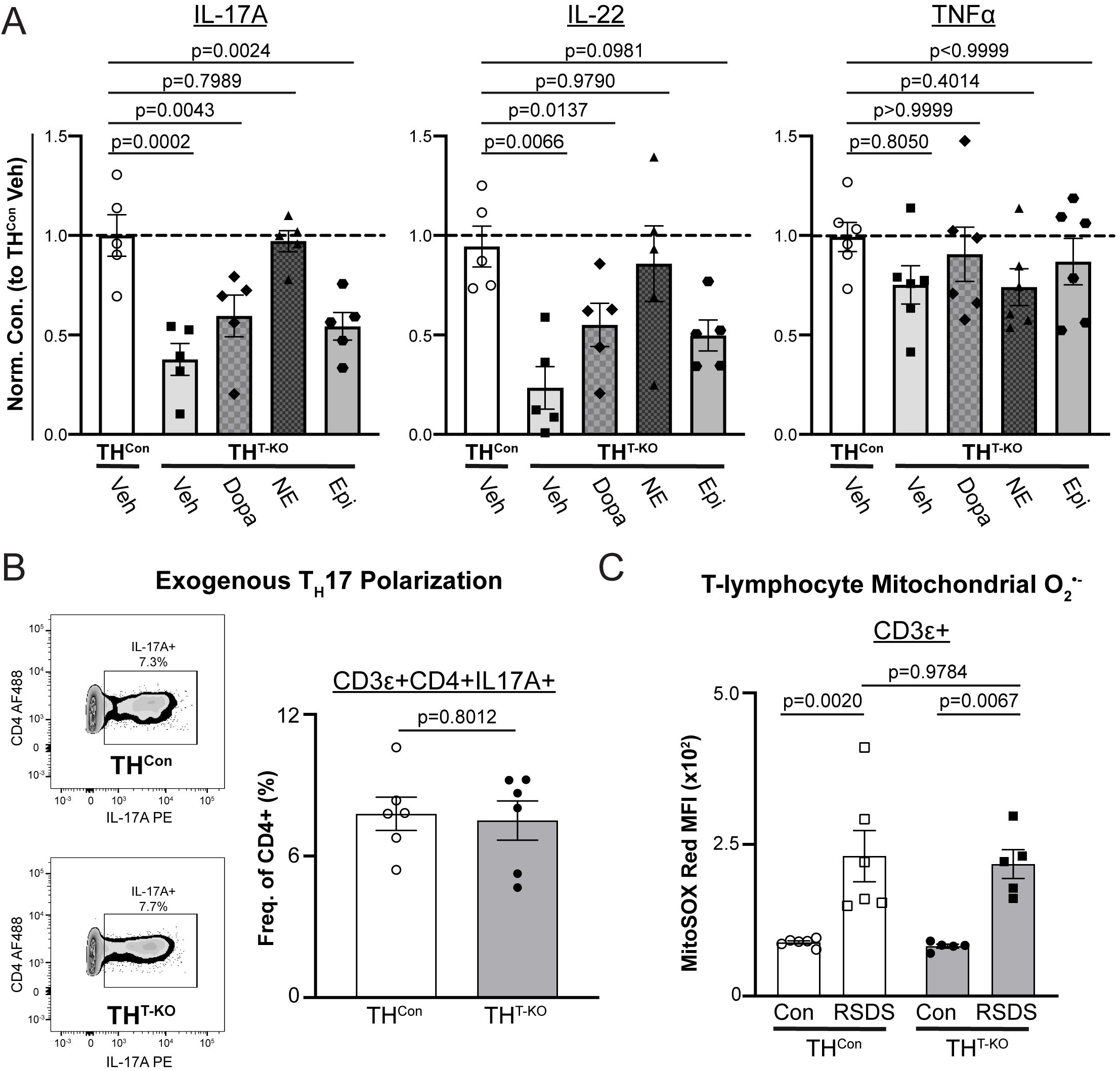
T_H_17 cytokine production can be rescued with catecholamine supplementation, but T-lymphocyte TH is not necessary for T_H_17 polarization, nor does it rely on mitochondrial redox. A) CD4+ T-lymphocytes were isolated and activated with 10 μM of respective catecholamines supplemented. After 72 hours, spent culture media was collected and assessed for cytokines by Meso Scale Discovery T_H_17 (Combo 2) assay. Cytokine concentrations are normalized to TH^Con^ and final cell counts. P-values represent nonparametric, paired measures ANOVA multiple comparisons tests (Friedman) compared to TH^Con^ vehicle. B) CD4+ T-lymphocytes were isolated, activated, and cultured under T_H_17 polarizing conditions for 5 days, then analyzed by flow cytometry. *Left,* Representative zebra plot. *Right,* TH17 T-lymphocytes following polarization; p-value by unpaired t-test. C) Following RSDS or control-housing exposure, splenic T-lymphocytes (CD3ε+) were stained with MitoSox Red to assess mitochondrial superoxide levels. Two-way ANOVA results, Stress p<0.0001, Genotype p=0.7096, Interaction p=0.8952.

### T-lymphocyte TH is not necessary for ex vivo T_H_17 polarization and does not function through a mitochondrial superoxide mechanism

We next investigated necessity versus sufficiency of T-lymphocyte TH in T_H_17 polarization *ex vivo.* CD4+ TH^Con^ and TH^T-KO^ T-lymphocytes were isolated and cultured with T_H_17 exogenous polarizing conditions. T_H_17 polarization was evident by flow cytometric analysis after 5 days, with no observable changes between genotypes (p=0.8012, **Figure 6B**), suggesting T-lymphocytes can still polarize to T_H_17 in the absence of TH if given a strong enough stimulus. Lastly, as we have previously shown a role for mitochondrial superoxide in T_H_17 polarization,^16,25, 26^ mitochondrial superoxide was assessed herein. Unexpectedly, mitochondrial superoxide was increased by RSDS in both TH^Con^ and TH^T-KO^ T-lymphocytes (p=0.0020, TH^Con^ Con vs. RSDS; p=0.0067 TH^T-KO^ Con vs. RSDS; **Figure 6C**), implying an uncoupling of between T-lymphocyte TH mechanisms and the mitochondrial redox environment.

## Discussion

Herein, we successfully generated a murine TH-deficient T-lymphocyte model in order to assess its role after psychological trauma. Firstly, we demonstrated the viability of TH^T-KO^ mice and a successful reduction in TH in TH^T-KO^ T-lymphocytes, while also finding patent RSDS-induced changes to anxiety-like or pro-social behavior in TH^T-KO^ mice. In circulation, RSDS exposure increased canonical pro-inflammatory cytokines in TH^Con^ animals, while TH^T-KO^ mice demonstrated attenuated IL-17A and IL-22. Additionally, we found reduced expression of these respective genes in isolated T-lymphocytes, in addition to reduced T_H_17 (CD3ε+CD4+CD8-IL-17A+) T-lymphocytes in RSDS-exposed TH^T-KO^ as compared to TH^Con^. When activated *ex vivo,* TH^T-KO^ CD4+ T-lymphocytes secreted less IL-17A and IL-22 in culture, with no growth disparities, which could be rescued with NE supplementation. Importantly, TH^T-KO^ T-lymphocytes were still able to polarize to T_H_17 under exogenous T_H_17 polarizing *ex vivo* conditions, and RSDS-induced mitochondrial superoxide shifts were not influenced by TH in T-lymphocytes. Overall, this investigation provides new insight into the role for TH in T-lymphocytes in the context of psychological trauma, while making way for further neuroimmune investigations.

The genesis for this investigation was based on a simple finding from our group^19^. Across animals in both RSDS and control conditions, the gene expression of IL-17A and TH was strongly, positively correlated (R=0.8722) in purified T-lymphocytes. Furthermore, RSDS increased expression of TH within T-lymphocytes. To test the potential causal relationship of these findings, we generated a conditional T-lymphocyte-specific knockout model of TH. While sparse, previous reports have identified catecholamine generation from T-lymphocytes, but have primarily focused on *in vitro,* pharmacological approaches. More recently, Yang *et al.* created a TH-deficient T-lymphocyte model using a ROR-γt promoter-driven cre recombinase after demonstrating a selective increase in the epinephrine and phenylethanolamine N-methyltransferase (PNMT; the synthetic enzyme preceding its production) in T_H_17 polarized cells^28^. After deleting TH in T-lymphocytes, they examined the phenotype of experimental autoimmune encephalitis (EAE), but interestingly did not observe any difference in clinical scores or lymphocyte infiltration into the CNS. Critically, EAE is an antigen-dependent response that develops after immunization by myelin basic protein, whereas RSDS-induced pro-inflammation and IL-17A is not known to be mediated through a single specific antigen, and may be a much weaker immune stimulus compared to an autoantigen. Additionally, much of the work by Yang *et al.* assessed the role of TH in T-lymphocytes through use of *in vitro* assays where T_H_17 polarization was accomplished by exogenous administration of polarizing cytokines *(i.e.,* anti-IFN-γ, anti-IL-4, IL-6, TGFδ). In the work herein, we also demonstrate that T-lymphocyte TH is not necessary for exogenous, *ex vivo* T_H_17 polarization. Together, our works provide valuable insights into the critical relationship between T-lymphocyte-derived catecholamines and T_H_17 T-lymphocytes, with two differing disease models and approaches showing differential effects of TH loss.

Importantly, the exact mechanism that connects T-lymphocyte-generated catecholamines and T_H_17 cells is still unknown. In our previous work, we have demonstrated the role for neuronally-derived NE in driving T_H_17 profiles through a mitochondrial superoxide mechanism^25^. However, the data presented herein indicates that T-lymphocyte mitochondrial superoxide operates upstream or in parallel of T-lymphocyte TH in inducing T_H_17, since it was increased after RSDS independent of genotype. The loss of TH within T-lymphocytes would result in deficient production of all catecholamines, which canonically bind adrenergic receptors to influence T-lymphocyte functionality. There is a breadth of literature which has focused on the intracellular cascade that follows adrenergic receptor (AR) binding, and there are several potential pathways which could explain this relationship. In our work herein, we were able to rescue the attenuated *ex vivo* production of IL-17A and IL-22 by TH^T-KO^ T-lymphocytes through primarily norepinephrine supplementation, thereby further validating their role in this overall mechanism. While outside of the scope of this work, future work could serve to utilize various adrenergic α and δ agonists and antagonists to further refine this pathway. Additionally, delineating the reception of T-lymphocyte-generated catecholamines specifically is an important future direction, especially giving special attention to non-canonical reception of catecholamines and intracellular signaling, as reviewed thoughtfully by Bellinger *et al^29^.* It should be noted that adrenergic pharmacological activation and inhibition of T-lymphocytes presents a number of technical challenges, further discussed in our previous work.^15^

This work is not without limitations. RSDS is murine model of PTSD which induces a robust T-lymphocyte inflammatory response.^19, 24, 25^ However, RSDS utilizes retired male breeder CD1 mice to induce psychological distress, and thus limited the inclusion of female mice herein. While models of RSDS have been developed which utilize females, many of these often utilize differential stress induction for the female mice, which introduces further variability and precludes direct comparison between male and female mice. As new PTSD models adjacent to RSDS are further developed and refined^30^, examining the phenotype in TH^T-KO^ is an important extension of these investigations. Additionally, cre recombinase in our experimentation was driven by the distal promoter of Lck in our model. This cre activation at the double positive (CD4+CD8+) thymocyte stage results in cre expression in all αδ T-lymphocytes^23^. Since this is relatively early in T-lymphocyte development, TH knockout at this stage (and subsequent T_H_17 dysregulation) could be the result of a developmental defect. Additionally, IL-17 and IL-22 are also produced by γδ T-lymphocytes, which additionally can utilize the distal promoter of Lck^31^, thus indicating their possible contribution to the phenotype seen. Future work could further describe the involvement of γδ T-lymphocytes in pro-inflammatory cytokine production after psychological trauma exposure.

Overall, this work provides new insights into the role for T-lymphocyte TH, specifically during psychological trauma. By utilizing an *in vivo* model, we were able to effectively demonstrate how T-lymphocyte-generated catecholamines are necessary for the T_H_17-skewed inflammation seen during RSDS. Continued work investigating the mechanism of T-lymphocyte generated catecholamines, as well as their potential clinical relevance is an important future direction. By fully elucidating the nuance of these neuroimmune connections, we can further our understanding of fundamental T-lymphocyte biology and inflammation associated with PTSD.

## Acknowledgements

This work was supported by the National Institutes of Health (NIH) R01HL158521 (AJC) and F30HL154535 (SKE).

## Author Contributions

SKE and AJC conceptualized the overall investigation. SKE, CMM, GFW, and AJC designed all research methods and experimental studies. SKE, CMM, and GFW conducted experiments and analyzed data. SKE and AJC wrote the manuscript. All authors reviewed, edited, and approved the manuscript. AJC provided primary experimental oversight.

